# A model-centered introduction to curiosity

**DOI:** 10.1101/2025.11.19.689250

**Authors:** Augustin Chartouny, Benoît Girard, Mehdi Khamassi

**Author notes:** equal contribution (Benoît Girard, Mehdi Khamassi).

## Abstract

Curiosity, defined as a transient motivational state for information, explains many non-greedy decisions in humans. Novel, surprising and uncertain situations are thought to elicit curiosity, leading to various models for information-seeking behaviors. We present a unified comparison of curiosity models based on experimental results and previous models from the decision-making and reinforcement learning literature. We provide and explain at least one mathematical formula for each curiosity category, implement them, and compare them on the same uncertain and changing task. This comparison illustrates how motivation and exploration may evolve differently based on what elicits curiosity. This work promises to increase the understanding of curiosity models and provide a synthesized toolbox for future models of information-seeking behaviors in animals or artificial agents.

## 1 Introduction

### 1.1 What is curiosity?

When an animal explores unknown surroundings, it engages in a potentially dangerous and energy costly activity. Yet, animals and humans sometimes preferentially explore novel, surprising, or unknown situations over familiar ones. For example, rats spontaneously explore novel mazes [1], and preferentially explore novel mazes over known ones [2]. The goal of this spontaneous exploration may be to find resources, to make sure that the environment is safe, to become better at predicting and controlling the environment, or simply to avoid boredom and remain aroused [3].

Curiosity has been introduced to explain this spontaneous exploration as a transient motivational state for information acquisition [3, 4, 5, 6]. As such, curiosity has been found to bias humans’ decisions towards informative options, while improving learning and retention of information [7]. This broad definition of curiosity as a transient motivational state for information leads to concurring characterizations of curiosity. Curiosity has been described variously as a drive [3], a desire, or a transient mental state [4], that can be experienced as positive (interest) or negative (deprivation) [8], and is typically intrinsic but can be modulated by extrinsic incentives [5]. In particular, intrinsic motivation and curiosity are often used interchangeably in the machine learning literature [9].

Prior work studied the influence of curiosity on animal and human decision-making [6]. Curiosity has been found to bias human and animal decisions towards informative options, resulting in goal-directed exploration. For example, humans explore preferentially options on which they have little information [10], which are of intermediate difficulty [11], or which are novel or surprising [12]. Importantly, children and various animal species exhibit information-seeking behaviors [13], showing that curiosity may be a key component of animal cognition. Recent literature also demonstrated that curiosity increased learning, even when controlling for potential confounding factors such as prior knowledge [14]. For example, humans remember better the answer to a trivia question they were curious about [7] and 11-month infants explore preferentially and remember better what violates their previous knowledge [15]. A potential hypothesis to explain increased learning is that people engage better when they are curious, leading to improved attention and learning. Hence, curiosity has been found to increase exploration and learning simultaneously.

Most studies indicate that curiosity primarily directs exploration and improves learning towards what initially sparked it [16]. However, curiosity can also improve the learning of incidental items. In a trivia task, human faces were presented to human participants between a question and its answer. Participants remembered faces better when the question had elicited high curiosity [17], as long as these faces appeared soon after curiosity was sparked [18]. In a similar paradigm with scholastic facts instead of faces, curiosity impaired retention of incidental information, likely due to competition for cognitive resources [19]. Overall, these results suggest that curiosity mainly promotes directed exploration and learning, which may spill over to support incidental learning or undirected exploration.

Based on the nature of the behavioral tasks, curiosity has been associated with various brain regions [20]. Anterior regions of the prefrontal cortex in humans and non-human primates have been associated with information-seeking behaviors [21, 22]. The ventromedial prefrontal cortex, the midbrain dopamine neurons, and the ventral striatum have been found to concurrently encode subjective values and information in humans and non-human primates [23, 24]. These results support a common-currency hypothesis for information and rewards provided externally by the environment or the experimenter, such as food or money [25, 26]. Finally, the hippocampus has been more activated in curious trials than non-curious trials, supporting the role of curiosity in learning and memory formation [7, 17].

Depicting curiosity as a transient motivational state for information separates the state of curiosity to its causes and consequences (Figure 1). With this framework, curiosity is not an umbrella term for its causes, or directly identified with increased exploration and learning. This distinction highlights that the causes and consequences of curiosity may vary based on confounding factors, such as arousal, stress, or noise in the learning process.

**Figure 1.**
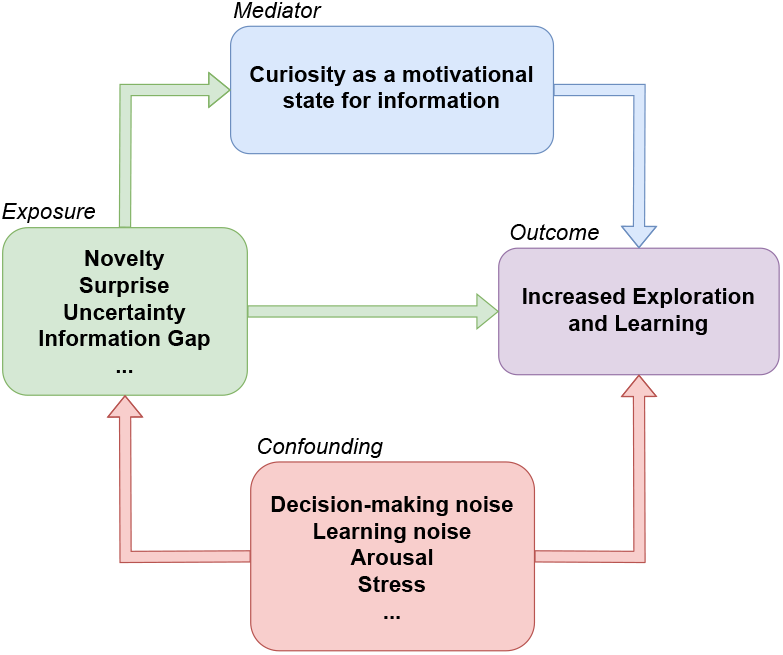
Curiosity as a motivational state for information. Being exposed to novel, surprising, or uncertain events may favor exploration and increase learning. This phenomenon may be partially mediated by curiosity, defined as a motivational state for information. Exploration, learning, and situations which elicit curiosity are influenced by other factors such as stress, arousal, or noise.

### 1.2 Choice and learning variability beyond curiosity

Choice and learning variability in humans do not only stem from curiosity or external incentives such as food or money. For example, stress and anxiety may reduce exploration [27], while positive emotions may facilitate learning [28]. Importantly, some factors, such as attention or effort, may influence exploration and learning, but also the perception of what is novel, surprising, or uncertain [29].

Additionally, noise in the learning or decision-making process may explain choice and learning variability in humans and animals [30, 31]. Noise-based exploration is typically associated with undirected exploration, while curiosity is often associated with directed exploration [32]. The distinction between undirected and directed exploration in humans has been emphasized by the horizon task [10, 22]. In this two-armed bandit task, participants must maximize their gains by accumulating rewards from two slot machines. They have partial information about the reward distributions of the machines, and a limited horizon to maximize their gains: either one or six decisions. In this behavioral task, exploration is said to be undirected when participants have an equal amount of information between the two machines but choose the one with the smallest expected gain. Contrarily, exploration is said to be directed when participants select the bandit arm with the smallest expected gain and the least information (and thus the highest potential to gain information). Although the term *random* is sometimes used to describe undirected exploration [10], this terminology may be confusing: undirected exploration does not always correspond to a random choice over the action space.

A common approach to model undirected exploration is to use a function modeling variability at the decision-level directly such as the softmax function [33, 34]. Using the framework of Markov Decision Processes, the probability of choosing an action *a* in a state *s* depends on the expected utility *Q*(*s, a*) (named Q-value) of this action (Equation 1). Softmax action selection reads

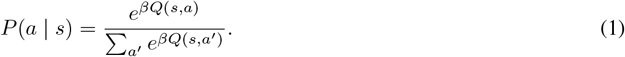

If all Q-values are null, softmax action selection is random over all actions. Otherwise, it is biased towards higher Q-values, with an inverse temperature parameter *β >* 0. The smaller the *β*, the more the agent chooses non-greedy actions (actions with *Q*(*s, a*) *<* max_*a*_*′ Q*(*s, a*^′^)). This formulation for undirected exploration has been widely used to model human behavior or to induce simple exploration in artificial agents [35, 36, 37]. The value of the inverse temperature parameter *β* in the softmax decision-making process can be adapted online, for example, based on the performance of the agent [38].

Another method for undirected exploration, named *ϵ*-greedy, consists in choosing a random action with a 0 *< ϵ <* 1 chance, and the best action with a 1 − *ϵ* chance. When *ε* is fixed, this action selection method grants an irreducible exploration rate and may offer better theoretical guarantees than softmax [39]. However, *ϵ*-greedy assumes that agents can choose actions randomly, and does strong predictions about decisions at each step (with a 1 − *ϵ* probability). This often results in much lower scores of performance than softmax when trying to explain human behavior using classical information criteria [40].

Methods such as *ϵ*-greedy and softmax model choice variability at the decision-level. However, choice variability may come from noise in the learning process instead [31]. A common way to model errors in the learning process of value-based tasks is to add random noise when updating the Q-values [36]. With a classical model-free update and an *α* learning rate,

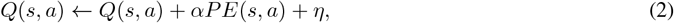

where *η* is a noise term sampled from a noise distribution at each update, and *PE*(*s, a*) is the prediction error the agent has just gotten when performing action *a* in state *s*. The noise term *η* can follow a uniform or Gaussian noise function, or increase with the size of the update [41]. The value-based computation from Equation 2 can be extended to non-value-based learning. For example, learning noise can be added to a probability distribution every time it is updated.

These results indicate that behavioral variability rely simultaneously on information-seeking and decision-making or learning noise. Many models of human behavior or autonomous learning combine undirected exploration, coming from noise in the learning or decision-making process, with directed exploration, coming from overt information-seeking [42]. Similarly, in our simulations, we use information bonuses to direct exploration and softmax decision-making (Equation 1) for undirected exploration and choice variability.

### 1.3 Situations which elicit curiosity

Curiosity directs exploration and evolves over time differently based on what initially sparked it. Previous work highlight that novelty [43], surprise [12], uncertainty [44], intermediate complexity [11], and a few other situations may elicit curiosity [6]. These various situations lead to different models, such as novelty-driven, surprise-driven, or uncertainty-driven curiosity [12, 45, 46]. The main goal of this article is to describe how each situation may induce curiosity and drive its evolution over time. To compare situation-driven models of curiosity, we provide one mathematical model for each candidate situation and simulate them on the same task. To direct exploration, we use information bonuses, often referred to as intrinsic rewards in the machine learning literature [9]. Information bonuses are summed by the agent with explicit rewards from the environment, following *R* = *R*^Bonus^ + *R*^Env^. This classical hypothesis is consistent with many findings in human and animal behavior indicating that information bonuses are comparable to rewards [47, 20].

## 2 Information-driven models

The information-seeking models we present apply to any discrete Markov Decision Processes. This framework encompasses many decision-making tasks such as bandit tasks or two-step tasks [48, 49]. The models we describe refer to an agent, *i*.*e*., an animal, a human, or an artificial decision-maker, interacting with an environment. The agent makes an action *a* in a state *s*, leading to the observation of an arrival state *s*^′^ and a potential reward *r*. The expected utility of each action *Q*(*s, a*) is a function of the predicted distribution of observations *P* (. | *s, a*) and the expected average reward *R*(*s, a*) the agent learns online. We denote *n*(*s, a*) the number of times the agent performed action *a* in *s, P*_*n*_(.| *s, a*) and *R*_*n*_(*s, a*) the associated models of the agent formed with *n*(*s, a*) observations, and *s*^′^ the observation at the *n*(*s, a*)-th choice. Further information about how agents may learn *P* and *R* are provided in the detailed methods (section A).

### 2.1 Novelty

Novelty-driven curiosity biases the agent’s behavior towards unexplored actions or unknown outcomes. The most common method to model novelty-seeking methods is to add an optimistic bonus in the face of novelty [50, 51, 52]. Consequently, the information bonuses of novelty methods decrease over time, as the agent repeats decisions or encounters previously observed outcomes. Novelty-based methods are often used in large or continuous environments, either to improve the performance of artificial agents [53], or to explain human behavior [43]. To categorize novelty methods, we make an explicit distinction between novelty in the behavioral space (have I ever done this action before?) and in the observational space (have I ever witnessed this outcome?). We call these methods action novelty and state novelty.

#### 2.1.1 Action Novelty

Action-novelty-seeking methods promote a transient initial exploration in new environments. For example, optimistic initialization of the Q-values in the reinforcement learning framework is a very common method to bias exploration towards new actions [39]. In decision-making and in reinforcement learning, many models are optimistic in the face of action novelty. For example, the model explaining the participants’ behavior in the original version of the horizon task is optimistic in the face of novelty [10]. It uses the difference between the number of times each of the two bandit arms have been selected as an information bonus. Similar count-based methods, which link exploration to the number of times one action is chosen, have been extensively studied in reinforcement learning. For example, the information bonus can decrease linearly with the number of times one action has been selected [52], with a square root [50, 54] or be constant for a predefined number of passages before being set to zero [51]. This information bonus generally reads

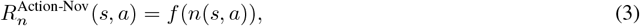

where *f* is a positive function decreasing with the number of times *n*(*s, a*) the agent chose action *a* in state *s*, and where 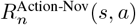 is the action-novelty bonus for (*s, a*) once the agent already chose *n* times action *a* in *s*. One limitation of count-based methods on the number of actions is that they assume that all actions provide the same amount of information, irrespective of their outcomes. Hence, they fail to capture the capacity of humans to generalize the outcomes of similar actions [55], or the potential interest of animals for actions with uncertain outcomes over predictable ones [56]. These shortcomings are compensated by the simplicity of action-novelty methods, which are easy to compute and do not rely on any feedback from the observational space.

#### 2.1.2 State Novelty

State novelty, defined as novelty in the observational space, often requires further computation than action novelty. For example, state novelty can be measured with a distance assessing how novel an observation is as opposed to previous ones [57]. Hence, state novelty necessitates predictions about the potential observations, or prior knowledge about the environment. Additionally, novel observations may elicit surprise and novelty at the same time as they are not expected by the agent’s model [12]. To differentiate state novelty from surprise, state novelty is often described as a measure of unfamiliarity, while surprise is described as a measure of unexpectedness [58, 46]. This distinction indicates that an observation is novel if it has not been experienced many times, irrespective of the action or the situation that led to it. On the contrary, surprise is context-dependent, as it relies on the observation prediction for the current action in the current state [46].

Similar to action novelty, the novelty of an observation decreases with the number of times it has been observed. Hence, to model state novelty, actions which tend to lead to new observations must provide a larger information bonus than the ones leading to known outcomes. With *N* (*s*^′′^) the total number of times the agent observed a state *s*^′′^, the model for state-novelty action we introduce reads

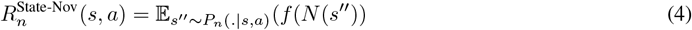

where *f* is a positive function decreasing with the total number of times *N* (*s*^′′^) the agent observed *s*^′′^. Equation 4 is the expectation of witnessing novel observations based on the current model of the agent. At start, the agent may have little prior information on the potential observations it can encounter. This means that action novelty may lead behavior before state novelty, and that combining both novelty types may be key to capture the behavior of an agent exploring a novel environment.

### 2.2 Surprise

Surprise is a transient response to an unexpected event, and is often explained as a prediction error coming from a mismatch between the agent’s internal model and an observation [12]. Surprising events have been shown to motivate exploration in humans. For example, infants and children explore and learn more effectively from objects which violate their expectations [59, 15]. Several physiological signals have been associated with surprise in humans, such as the P300 signal in event-related potentials, or an increase in pupil size [60, 61]. A recent work tried to categorize surprise definitions based on their formulas, showing that most formulations were functions of predictive or update surprise [62]. Predictive surprise is linked to the likelihood of observing an event under a model [63]. Conversely, update surprise is linked to the size of the update in the model after observing an event [64]. Hence, update surprise represents a relative surprise corresponding to an update in the model (does my model change after observing this event?) whereas predictive surprise represents an absolute surprise (is this event unexpected by my model?).

One of the main differences between update and predictive surprise is that they do not predict the same behavior when there is irreducible noise in the observation distribution. For example, predictive surprise predicts that agents remain interested in a white noise TV, although the signal is unpredictable. Contrarily, update surprise eventually decreases after observing that the white noise is unpredictable and that the learned model does not improve [64]. Predictive surprise and update surprise have been associated with different brain regions [61, 65].

#### 2.2.1 Predictive surprise

Predictive surprise is often modeled by reward or state prediction errors in decision-making tasks [49]. In value-based tasks, reward prediction error can be modeled with the value error coming from TD-learning-based algorithms [66]. State prediction error can be modeled using Shannon surprise, as the log-likelihood of observing an event under a model [63]. When using Shannon formula, predictive surprise reads

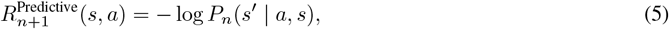

where *s*^′^ is the observation from the agent the *n*(*s, a*)-th time it took action *a* in *s*. Hence, at each time-step, the surprise bonus associated to (*s, a*) from Equation 5 relies on the observation at the previous passage. In short, Shannon surprise used as a drive for information favors actions which have just led to a surprising observation. To motivate exploration while not relying on the last surprising observation only, one can smooth predictive surprise over several trials.

#### 2.2.2 Update surprise

A second aspect of surprise that has been well-studied in the literature is surprise linked to belief updates, sometimes coined Bayesian surprise in the Bayesian inference literature. It refers to how much one observation improves the internal model of the agent [64]. Update surprise identifies better to an update signal related to information gain than to a surprise signal. For example, predictive surprise explains the P300 component of the human event-related brain potential better than update surprise [60]. Similarly, predictive surprise has been found to correlate positively with pupil size while update surprise decreases with it [61]. Hence, surprise may not be the best-suited terminology to describe this information bonus.

A possible expression for update surprise in Markov Decision Processes reads

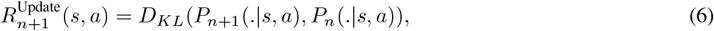

where *D*_*KL*_ is a Kullback-Leibler (KL) divergence. In Equation 6, 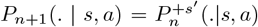 is the probability distribution built on the prior distribution *P*_*n*_(*s, a*) and the arrival state *s*^′^ observed at the *n*(*s, a*)-th passage. Thus, update surprise depends directly on the most recent observation and on the update rule for *P*. Equation 6 is a simplified version of Bayesian surprise with directly observable variables [64].

Several formulations of update or Bayesian surprise invert the order of the two distributions in the Kullback-Leibler (KL) divergence [62, 67, 68, 60]. We found no explanation for this inversion in the literature. This inversion may facilitate computations, as the support of the posterior distribution should contain the one of the prior distribution. However, it does not account for surprising events well, as the divergence does not diverge when the agent observes something completely unpredicted. In our simulations, we stick to the posterior-prior order as it seems to represent a surprise signal best (see Figure S6 for a comparison).

### 2.3 Uncertainty

Uncertainty is linked to the perceived level of variability and noise in the environment [35]. From the agent’s perspective, uncertainty is often categorized using the expected-unexpected dichotomy [69]. Expected uncertainty is the uncertainty predicted by the model that cannot be reduced, which may be due to the inherent stochasticity of the environment. In contrast, unexpected uncertainty is the uncertainty that was not predicted by the model and that can and should be reduced to improve predictions. Expected and unexpected uncertainties can be formulated using averaged versions of the surprise formulas we presented in Equations 5 and 6.

#### 2.3.1 Expected uncertainty

Previous work used the variance of the internal model to represent expected uncertainty [70, 71]. For example, a variance-based upper-bound confidence approach favors choices with high uncertainty [72]. Another model, named Thompson sampling, increases choice variability based on the total uncertainty on the environment, irrespective of where this uncertainty comes from. Thompson sampling approach is similar to modifying the *β* parameter in the softmax formula based on the expected uncertainty (Equation 1). A more general approach consists in using the entropy of the distributions instead of their variances. To compute the entropy on the observations for one action, we take the expectation over the predictive surprise (Equation 5). The resulting information bonus reads

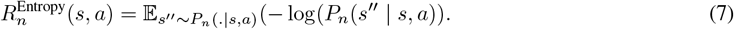

The entropy formulation from Equation 7 is prospective, while predictive and update surprises are retrospective. Taking variance or entropy directly as an information bonus motivates agents to explore areas they are uncertain about. However, in stochastic environments, this bonus does not completely vanish over time, even after learning, as environmental uncertainty is irreducible. Thus, expected-uncertainty-driven agents always prefer unpredictable actions over predictable ones. Expected uncertainty can be combined with novelty methods to reduce the information bonus as the agent samples more observations (Figure 4, top).

#### 2.3.2 Unexpected uncertainty

While expected uncertainty represents the agent’s expectation on the inherent uncertainty of the environment, unexpected uncertainty refers to an unexpected and subjective change perceived by the agent [71]. Hence, unexpected uncertainty is often associated to strong or fast belief updates in volatile environments [35]. Contrary to expected uncertainty which is purely prospective, unexpected uncertainty accounts for surprising events leading to belief updates. Thus, to model unexpected uncertainty, we temporally extend the formulation of update surprise from Equation 6. Instead of computing the update between two successive passages, we compute the KL divergence between the current model and the same model without the *k >* 1 most recent observations.

#### 2.3.3 Expected information gain

To reduce unexpected uncertainty, we integrate update surprise from Equation 6. This information bonus is the expected information gain in the model after performing an action. Expected information gain reads

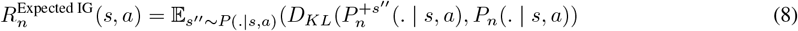

Expected information gain from Equation 8 represents how much the agent may learn by performing action *a* in state *s*. This bonus is an expectation over all potential model updates the agent may make based on the upcoming observation. This information bonus is prospective (how much information the agent may obtain), while update surprise is retrospective (how much information the agent obtained). Expected information gain naturally decreases when the agent becomes more certain about their model. In practice, computing this expected information gain may be computationally expensive, as it requires to build several distributions based on the potential future observations.

### 2.4 Learning progress

Learning progress is an information gain measure which favors exploration towards areas the agent can learn the most from [73]. This simple heuristic explains well how humans and artificial agents can learn tasks of increasing difficulty by monitoring their progress [74, 11]. However, monitoring learning progress is often task-dependent, leading to various task-specific formulas for learning progress [75]. Additionally, learning progress is used both in skill learning (variation in competence) and knowledge acquisition (variation in prediction errors). For example, based on the task to solve, a good proxy for learning progress can be the variation in the percentage of correct answers of a human participant, or the progress in the ability to reach different states of the environment for an artificial agent [76, 77]. This distinction between skill learning and knowledge acquisition is sometimes overlooked, leading to some misconceptions about learning progress, such as its role in goal-oriented behaviors [78]. In the following, we focus on learning progress as knowledge acquisition.

The seminal work which introduced learning progress used a difference of averaged prediction errors between two successive timesteps [73]. This formulation is consistent with progress in learning: if the prediction error of the agent decreases, the agent improves its model of the environment and is thus learning. Conversely, if the difference in prediction errors is null, the agent is not learning anymore. However, with this original formulation, increases in prediction errors, which notably happen when the agent faces surprising events, lead to negative learning progress. Thus, the agent receives a penalty when choosing actions which have just become surprising. This is detrimental when trying to adapt to changing environments, can increase catastrophic forgetting in machine learning, and fails to account for the interest of humans in surprising events [74, 59]. Adding absolute values solves this limitation, making increases in prediction errors as important as decreases in prediction errors [78]. This motivates the agent to explore areas where prediction errors vary strongly, whether they increase or decrease. Yet, empirically estimated learning progress is a proxy for what the agent can truly learn. For example, learning progress agents may suffer from some inertia, as they must sample actions several times to estimate their learning progress.

To study learning progress for model learning, we use the absolute difference in consecutive entropy. This method is a simplified version of previous work in model-based reinforcement learning [79, 80] and the same formulation without absolute values has been used previously [81]. As the absolute variation in the average predictive power of the agent, this learning progress bonus reads

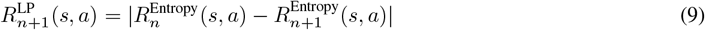

where 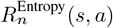 is the entropy of the predictive model of the agent after performing *n* times action *a* in *s* (Equation 7). To smooth learning progress over time, the agent can compare entropy between two non-successive passages, separated by *k >* 1 passages [79].

### 2.5 Model reliability

Humans and animals learn cognitive maps of their environment or the task at hand [82, 83]. Hence, model reliability, which represents the confidence of the agent in its internal model of the environment, may motivate exploration [64, 44]. For example, in uncued task-switching paradigms in humans, exploratory response rates increase a few trials after a change and decrease faster in recurrent tasks [84]. This potentially indicates that participants learn internal models and strategies for each task, and reduce their exploration rate as they become confident in their model of the task [85].

Some methods we previously discussed can account for model reliability. For example, update surprise assesses how much the model of the agent changes between two consecutive passages (Equation 6). A model that changes a lot based on incoming observations may not be reliable. Similarly, learning progress compares the predictive power of the model between two successive passages and can thus be identified as model reliability (Equation 9). Assessing model reliability on one observation only may lead to erroneous interpretations when witnessing very uncertain events. Thus, to measure model reliability, update surprise and learning progress can be computed over longer time windows.

Another option to measure model reliability is to directly compute how well the learned model explains recent observations [86]. Instead of measuring the variation in the model (update surprise) or the variation in the predictive power of the model (learning progress), we propose to measure how well the learned model explains recent observations. With *P*_*n*−*k*→*n*_(*s, a*) the model formed with the *k* most recent observations when performing action *a* in *s*,

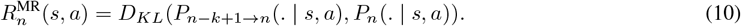

Model reliability compares the distribution of recent observations *P*_*n*−*k*+1→*n*_ with the learned model *P*_*n*_(*s, a*) (Equation 10). Using a KL divergence highlights that the two distributions are asymmetric, as the learned model has more observations and is potentially more precise than the model for recent observations.

## 3 Results

### 3.1 Experiments

We implement and compare all of the curiosity models we introduced (Equations 3, 4, 5, 6, 7, 8, 9, 10). For novelty agents, we use an information bonus inversely proportional to the square root of the counts, such that, for *n* the count, 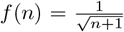 (Equations 3 and 4). We normalize the entropy and predictive surprise to compare them with the other methods (see Figure S5 for the plots without normalization). We present results for the *k*-step versions of update surprise, learning progress, expected information gain and model-reliability. For learning progress and update surprise, this consists in computing the update surprise or the learning progress between the current model and the same model without the last *k* passages (instead of making the computation between two successive passages). The *k*-step variation of expected information gain consists in computing how the model would evolve after witnessing *k* times one observation (instead of observing it once in the original formulation). We normalize expected information gain to compare it with other methods. The agents learn the probabilities over arrival states *P* (*s*^′^ | *s, a*) by computing the frequency over arrival states with a small uniform prior and a rolling window of *H* observations (section A).

To clearly identify the different properties of each curiosity method, we simulate a simple one-step environment. In this environment, the agent faces three dice of 6 faces numbered from 1 to 6. The first die is deterministic and always returns 1, the second one is slightly uncertain, returning 2 or 3 with a 50% chance, and the last one is fair, returning each value with a 1*/*6 probability. Importantly, after 100 choices, the distribution of the second die changes from 2 − 3 with a 50% chance each to 4 − 5 with a 50% chance each. This allows to study the evolution of curiosity in a simple, initially novel, uncertain, and changing environment. There is no external reward in the environment. Thus, the agents make decisions solely based on the information bonuses they compute. To choose actions based on their expected values *Q*(*s, a*) = *R*^*Bonus*^(*s, a*) and add choice variability, all agents use a softmax decision-making (Equation 1). We mainly describe the evolution of curiosity under two metrics: the value of information bonuses and the probability to choose each die (Figures 2 and 3). In all of our simulations and when not specifically indicating other values, we use *H* = 50, *β* = 10 (*β* = 3 for the entropy and the predictive surprise to add choice variability), and *k* = 10 for *k*-step variations of the curiosity methods. We evaluate the agents over 500 seeds and show the confidence intervals at 95%.

**Figure 2.**
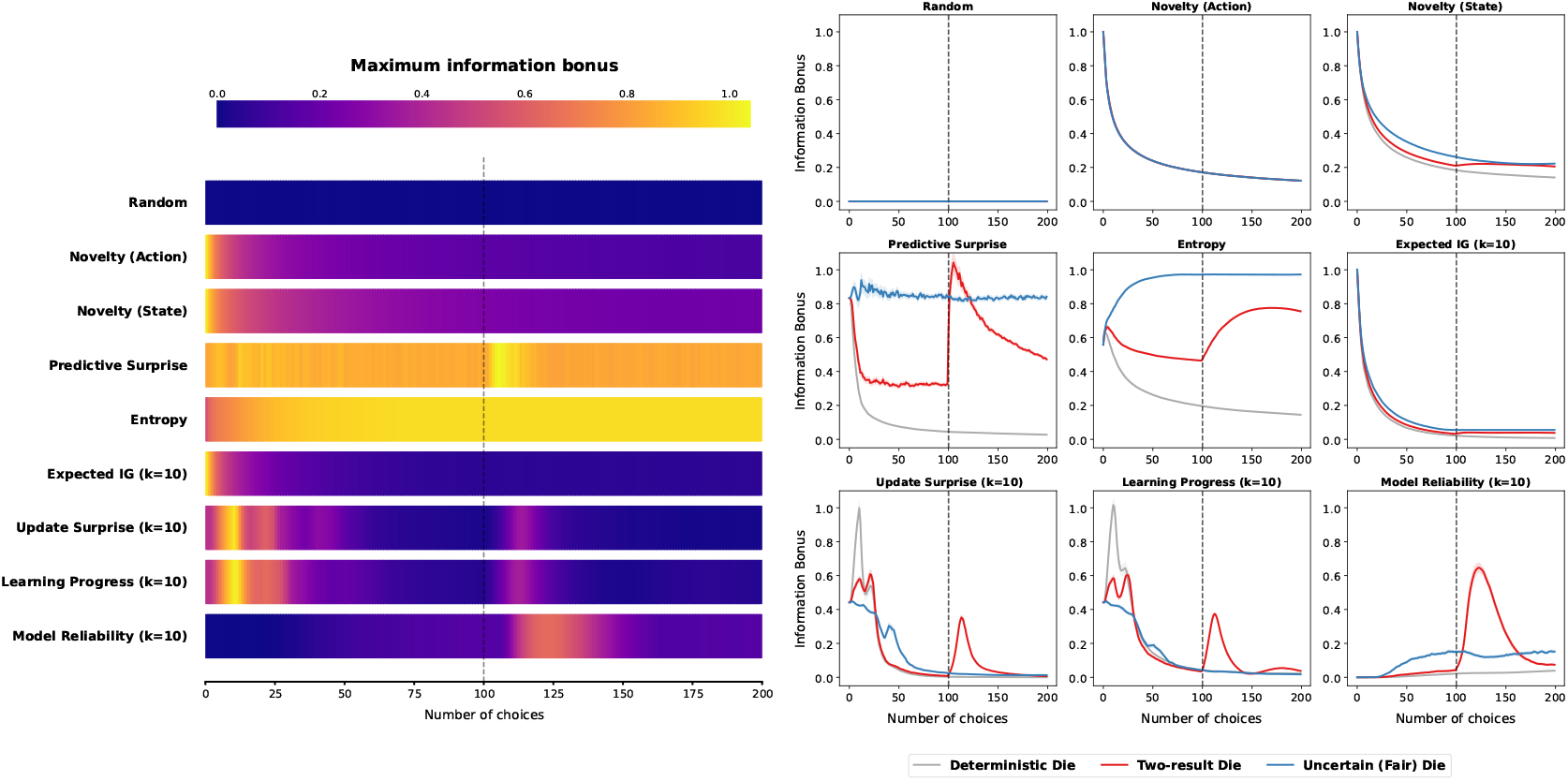
Evolution of the information bonuses over time for each curiosity method when facing three dice with varying uncertainty. At timestep 100, the outcome of the two-result die changes from 2 − 3 to 4 − 5 (with 50% chance each). This change is indicated with dotted lines. (Left) Comparison of the evolution of the maximal information bonus for each method over time. Most methods have high information bonuses at the start of the task and the die change induces an augmentation of the maximal information bonus for some of the curiosity methods. (Right) Individual evolutions of the information bonuses for each die and each method.

**Figure 3.**
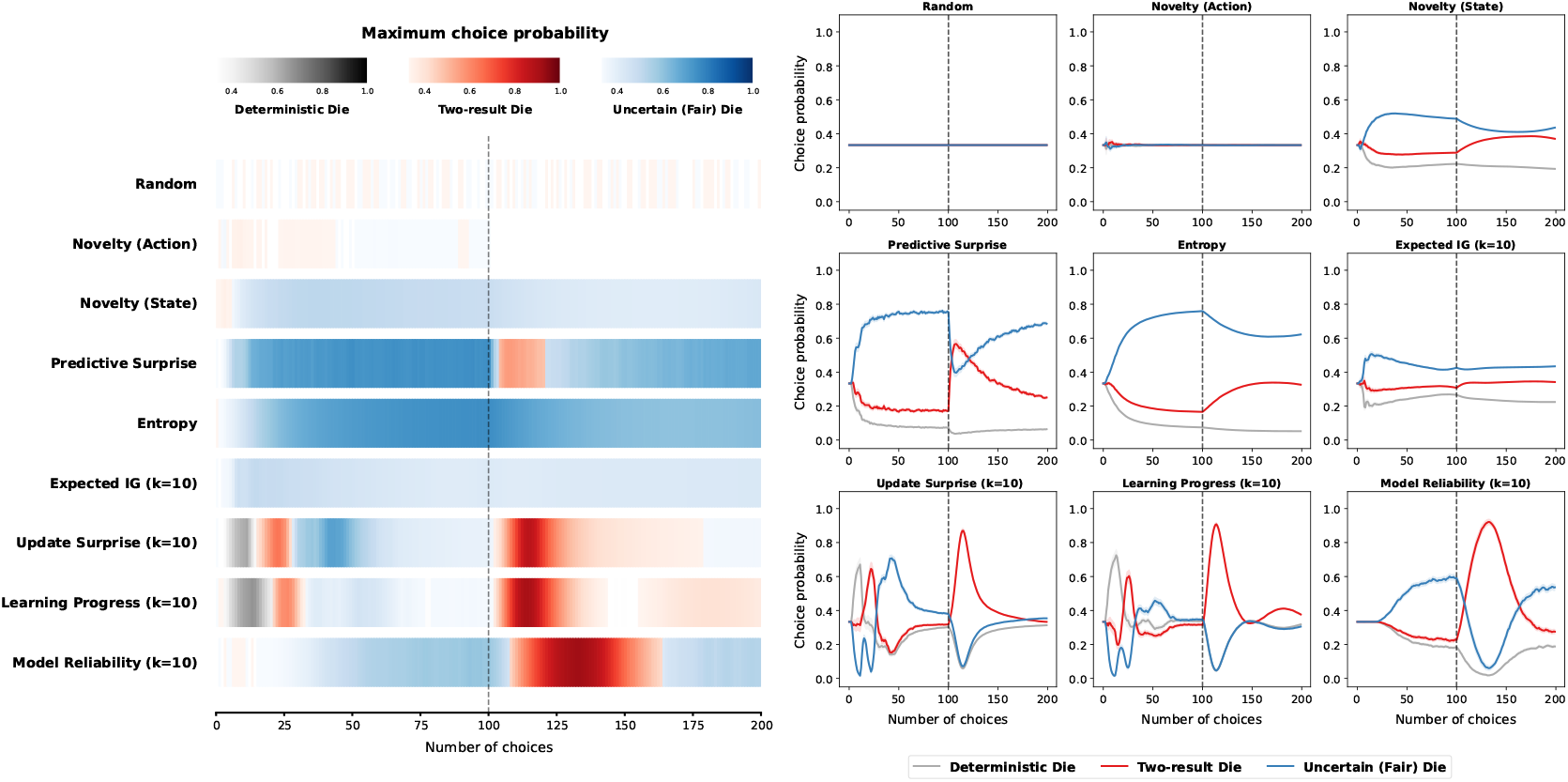
Evolution of the die preference for each curiosity-induced exploration method over time. At timestep 100, the outcome of the two-result die changes from 2 − 3 to 4 − 5 (with 50% chance each). This change is indicated with dotted lines. (Left) For each method, we indicate the die the agent preferentially chooses over time. The darker the color, the stronger the preference (in terms of choice probability). (Right) Detailed probabilities of choosing each die for each curiosity method.

**Figure 4.**
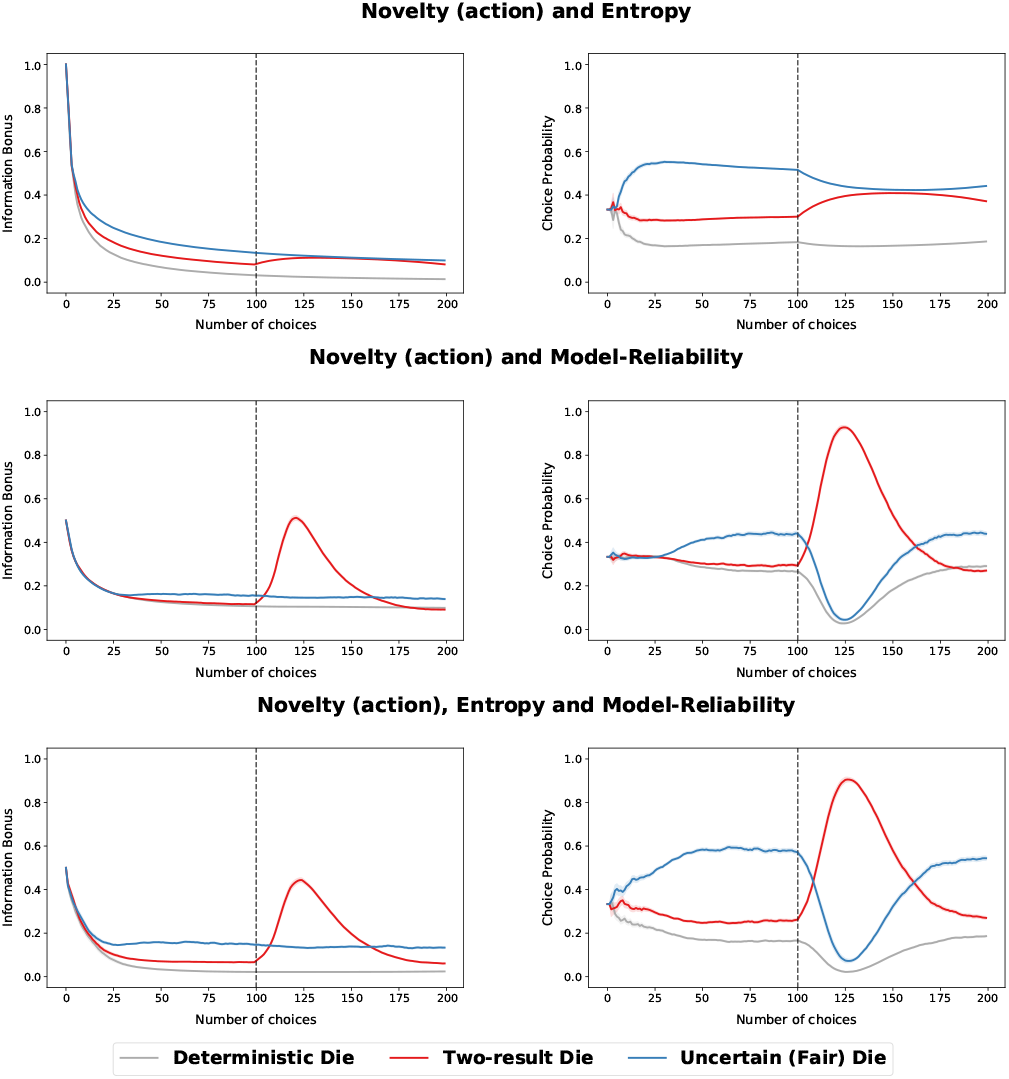
Evolution of information bonuses (Left) and choice preferences (Right) over time for the action-novelty and entropy (Top), the action-novelty and model reliability (Center), and the action-novelty, entropy, and model reliability (Bottom) information bonuses. Each plot is generated over 500 seeds and 200 trials. The two outcomes of the two-result die change at step *t* = 100.

### 3.2 Early exploration

Early exploration relies on what elicited curiosity, on the prior knowledge provided to the agent, and on the initial values set for the information bonuses. Importantly, the absence of prior knowledge on the potential outcomes, or a particular initial expectation on them, may lead to modified early exploration. In the dice experiment, we made the strong assumption that agents had an initial uniform prior for the dice outcomes, meaning that they knew that they were six-faced (from 1 to 6) and that all agents initially expected all dice to be fair (section A). We set initial information bonuses based on this hypothesis. Some methods do not need such detailed prior knowledge, such as the action novelty method which makes no assumption on potential observations.

In the very first steps and when averaged over 500 seeds, most agents have a close-to-random preference (Figure 3), coming from the initial uniform prior over the expected utilities of each action. After a few steps, some preferences emerge. The state novelty agent quickly prefers the action which grants the most novel observations, *i*.*e*., rolling the fair die. Similarly, entropy, predictive surprise, expected information gain and model reliability quickly favor the fair die over other options. This preference arises for different reasons based on the curiosity method utilized. The entropy method identifies the most uncertain distribution and prefers it strongly over the other ones. The predictive surprise approach fails to predict the outcome of the fair die and prefers it over other dice. Expected information gain identifies that it can learn more from the fair die. Finally, the dice models learned by the model reliability agent cannot explain perfectly most recent *k* = 10 observations when the dice have stochastic outcomes. Hence, the information bonuses for model reliability increase over time, with a preference for the uncertain die which provides the most variable observations. Action novelty shows little preference when averaged over 500 seeds. However, the high information bonuses at start indicate that action novelty has a more systematic early exploration than the random approach, by biasing the exploration towards actions that have not been tried enough. This systematic exploration brings convergence properties in multi-step Markov Decision Processes [54].

Two methods display very specific early exploration. The update surprise and the learning progress methods (with *k* = 10) show an early preference for the deterministic die first, the two-result die then, and finally the uncertain die. This behavior comes from the mismatch between the initial prior knowledge of the agents (that the dice are fair) and their observations. Thus, agents preferably sample from the distribution which is most different from their initial expectation (in terms of model update or entropy). Learning sequentially based on information gains is typical of curriculum learning methods [76].

### 3.3 Exploration in a stable environment

After enough observations, some agents tend to have a close to random preference, as their information bonuses strongly decreased over time. For example, learning progress and action novelty methods have no preference at the time of the die change (Figure 3). State novelty method has a small preference for the uncertain die before the change at *t* = 100. However, after more steps or by using a faster decreasing coefficient for the novelty methods (*i*.*e*., with a decrease of 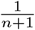), state novelty would have very small information bonuses and exhibit almost no preference. All information bonuses decrease over time but the ones from entropy, predictive surprise and model reliability (Figure 2). After enough observations, entropy and predictive surprise bonuses would eventually stabilize to values which reflect the inherent stochasticity of the dice.

### 3.4 Exploration when facing changes

Action novelty does not use any feedback from the environment and does not exhibit any change in die preference or maximal information bonus after the two-result die changes (Figures 3 and 2). Since the change appears to temporarily increase the number of observations of the two-result die from 2 to 4, some methods such as state novelty, expected information gain, and entropy slightly increase their information bonus for the two-result die when the task changes (Figure 2, Right). This change decreases the preference for the fair die, which remains a bigger attractor state for these methods, with higher entropy, number of observations, or expected information gain.

Update surprise and learning progress quickly focus on the changing die, as they detect a gap between what they know and what they observe. Update surprise increases because recent observations differ from what has been observed so far, modifying the model of the two-result die. The entropy of the two-result model temporarily increases as well, leading to a negative entropy difference and a positive absolute learning progress. Interestingly, learning progress shows a bi-phasic preference for the two-result die after the task change. This phenomenon, which is common in variation-based methods, is further described in the supplementary information (section C.2).

Model reliability detects that its model of the two-result die becomes unreliable, and directs its exploration towards it. This shift in preference is larger for model reliability than for learning progress and update surprise, but may seem to happen with a small delay (Figure 3, Left). When comparing the plots directly, the information bonus for model reliability increases as fast as the ones of learning progress and update surprise (Figure S2, Left). Notably, for all three methods, changes can only be detected when the agent performs the corresponding action. When controlling for the behavior of the agents at the time of the task change, the information bonus for model reliability increases as fast, if not quicker, than the ones from update surprise and learning progress (Figure S2, Right). These three methods can be used efficiently to redirect exploration after a task change, facilitating adaptation to the environment dynamics after a task change (Figure S1).

### 3.5 Combining approaches: Towards a unified model for curiosity

The formulas we presented are stereotypical as they each describe a specific situation which elicits curiosity. Models of curiosity which combine several approaches may thus explain animal behavior better than standard ones [72]. To illustrate how to build hybrid models, we mix action novelty with entropy, model reliability, or both. We introduce three corresponding new equations (Equations 11, 12, 13).

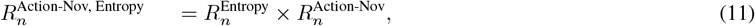

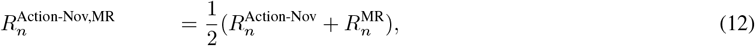

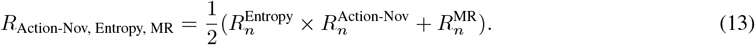

Multiplying entropy with action novelty (Equation 11) reduces the interest of the entropy method for the fair die over time, as larger entropy leads to more samples and thus less action novelty (Figure 4, top). This information bonus is more consistent with the hypothesis that humans may prefer slightly uncertain situations over fully predictable ones, and that this interest decreases when learners understand that the uncertainty is irreducible. This combined information bonus behaves closely to the state-novelty bonus in the dice experiment (Figures 2 and 3, Top-Right). However, the two approaches are theoretically distinct: the entropy computation does not use the nature of the observations while state novelty does. In our experiment, this results in the information bonus for the two-result die being larger for the action novelty and entropy method than for the state novelty method.

Summing action novelty with model reliability (Equation 12) combines early exploration from action novelty with the capacity to adapt to changes from model reliability. This provides an interesting information bonus which does need prior knowledge about the environment, is high when facing a novel environment, slightly prefers uncertain situations in stable environments, and increases when facing a change (Figure 4, center).

Adding a preference for uncertain situations on top of this bonus does not seem useful (Equation 13 and Figure 4, bottom), as the novelty action and model reliability agent already prefers uncertain situations in stable environments (Figure 4, center). Overall, combining information bonuses may elude some limitations of individual methods. For example, summing the action novelty and the model reliability bonuses endows model reliability with early exploration, or action novelty with change detection and a slight preference for uncertain situations in stable environments. This combined information bonus is a simple yet efficient model which may explain information-seeking behaviors in a wide range of experiments.

## 4 Discussion

We introduced and described several situations that may elicit curiosity. For each situation, we came up with at least one mathematical formula and studied how each formulation may be used to promote exploration in a simple three-action example. We highlighted how each method would promote early exploration, exploration in a stable environment, or re-exploration after a task change using the formalism of intrinsic reward in reinforcement learning [9]. Several results emerge from our simulations, such as the capacity of learning progress, update surprise, or model reliability to adapt to changing environments. We also illustrate how combining different information bonuses may improve information-seeking models. A few other factors that we could not implement in our framework such as goals, play, confidence, and empowerment, may participate in information-seeking behaviors or increased learning.

### 4.1 Other situations contributing to curiosity

#### Goals

Goals are central when studying curiosity, as curiosity is often directed towards a specific item, situation, or goal, and curiosity can help generate goals [73, 87]. For example, in tasks of curriculum learning, learning progress helps generating goals of increasing difficulty to direct exploration and learning [88, 89]. Several models of decision-making explore the link between goals and curiosity, either to explain human behavior [90] or facilitate skill learning in artificial agents [74]. Goal-conditioned approaches differ by what goals the agent generate and how they try to achieve them [91]. Thus, goal-based information-seeking methods tend to be task-specific. In the supplementary information, we present a simple goal-based approach with randomly generated goals (section C.6).

#### Confidence

Confidence refers to the agent’s subjective estimate of how likely their decision, judgment, or knowledge is correct or reliable. Confidence in knowing the answer to a trivia quiz question has been shown to vary in an inverted U-shape curve with curiosity [7]. However, the relationship between confidence and curiosity may vary based on the nature of the stimulus [92], or the wording of the question asking how confident participants are [14]. Several computational models of curiosity try to integrate a metrics for confidence in their equations [93, 62]. Confidence is a broad concept which partially overlaps with the uncertainty and model reliability formulations we presented.

#### Play

Play has been described as the will to seek out or create intricate and mildly complex situations, with the objective to resolve surprise and uncertainty [94]. Many theories describe play and its role in cognitive development [95]. Play stimulates curiosity, exploration and learning, having many potential functions, such as pleasure, practice, or performance [13]. Hence, framing tasks as games may benefit engagement, attention, or learning. However, we believe that providing a general formula to induce play-oriented behavior may be complicated, as play seems to rely on many mechanisms such as goal generation and surprise resolution, and is often a social mechanism involving several agents [13]. Thus, we could not come up with a task-general model for play.

#### Empowerment

Empowerment is a local state-based measure for controllability which consists in maximizing mutual information [96]. Empowerment favors states that offer a large number of outcomes and on which the agent has the most control. Empowerment is maximum when all actions have distinct deterministic outcomes, as the agent can reach many states and has a full control over the states it reaches. Importantly, empowerment measures potential controllability: the action distributions that achieve mutual information maximization may differ from the actual behavior policy [97]. One of the critiques of empowerment is that it is non-trivial to compute, as each computation requires solving a maximization problem over all possible action distributions [98, 99]. Empowerment has mainly been used in machine learning [100], but recent work suggest that it may be useful in explaining human exploration in open-ended tasks [101]. Empowerment-derived information bonuses are at the state level rather than at the state-action level. Hence, we could not implement empowerment in our framework.

### 4.2 Limitations and future work

Although we back up our model choices using literature in animal behavior and machine learning, some formulas we present may be debated. For example, our novelty models favor novel actions or observations. However, infants have been shown to prefer familiar stimuli over novel ones in various tasks [102]. Similarly, we provide one learning progress model for knowledge acquisition, but there exists several variations which may have been equally interesting [75].

In the supplementary information, we detail the methods we used and the results we found (sections A and B), and provide additional simulations and results for some of the approaches presented in the article (section C). Importantly, we highlight the limitations of some formulations we found in the literature. For example, the order of the two distributions in the update surprise computation matters (section C.4), and learning progress as a difference of prediction errors necessitates absolute values (section C.5).

Since this article shows how simple curiosity models drive information-seeking behaviors in a one-step task, future work could improve the models presented, apply them to a larger multi-step environment, or use them to explain animal data. For example, we did not implement increased learning based on the curiosity level [7, 103]. Future curiosity models could increase learning in curious trials by modulating the update of the learned model by the information bonus at the time of the observation. To apply these models to larger environments or animal data, one can use the formalism of Bayesian inference or model-based reinforcement learning (section A). We hope that the range of models we introduced or adapted from existing work, described, and compared, will help understand curiosity and its associated brain mechanisms.

## Acknowledgments

The authors would like to thank Jean-Pierre Nadal, Erik Nemeth, and Olivier Serris for discussions.

## Copyright

Article and figures by Chartouny, Girard, and Khamassi (2025); available under a CC-BY 4.0 license.

## Code

The code is available online on Github.

## A Detailed methods

### A.1 Explicit formulas used in the simulations

The formulas we present in the main article are general and can apply to many distributions or situations. In our experiment, we use discrete arrival states and actions which simplify the computations. Hence, equations 3, 4, 5, 6, 7, 8, 9, and 10 are implemented as equations 14, 15, 16, 17, 18, 19, 20, and 21 respectively.

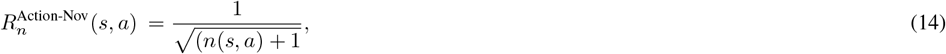

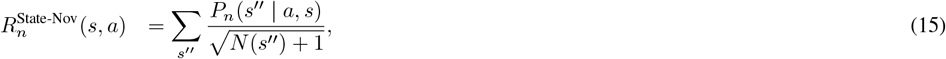

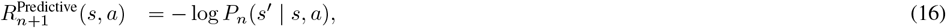

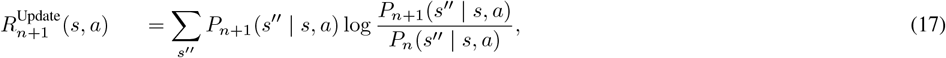

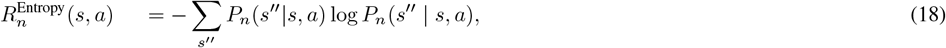

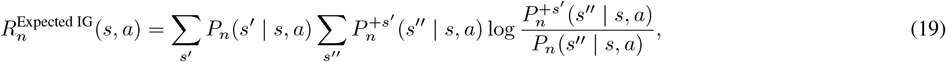

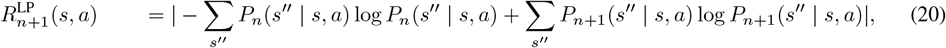

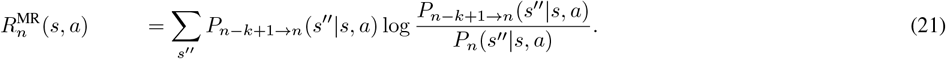

### A.2 Learning models

We compare different curiosity methods using the Markov Decision Process (𝒮, 𝒜, 𝒫,ℛ,) framework [39]. The agent chooses to carry action *a* ∈ 𝒜 in an environment composed of states *s* ∈.𝒮 When the agent takes action *a* in *s*, it arrives in a state *s*^′^ sampled from the transition function 𝒫 (*s, a*) and obtains a reward *r* sampled from the reward function ℛ (*s, a*). The agent estimates the transition 𝒫 and reward models ℛ of the environment using learning methods from model-based reinforcement learning.

Model-based reinforcement learning agents approximate the reward ℛ and transition 𝒫 models of the environment they interact with. The learning approach we implemented uses a frequentist finite-horizon approaches, forgetting oldest information when the number of observations in their model of (*s, a*) exceeds *H*. We denote *n*_*H*_ (*s, a, s*^′^) the number of times the agent reached *s*^′^ in the last *n*_*H*_ (*s, a*) ≤ *H* times it performed action *a* in *s*. Similarly, we denote *R*^*H*^ (*s, a*) the ensemble of rewards it received from the environment in its last *n*_*H*_ (*s, a*) passages. The approximate transition functions 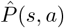 and reward function 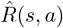 learned by the agent read

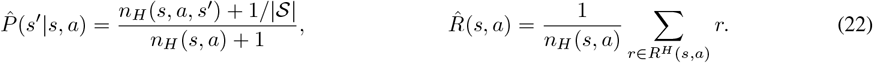

The approximate model of the transition function uses an additive smoothing factor of 1*/* |𝒮 |, with |*S* | the number of states in the environment (Equation 22). This smoothing factor indicates that the agent has an initial uniform expectation over the potential states it can reach. The agent adds internally generated information bonus *R*^Bonus^(*s, a*) with the reward provided by the environment 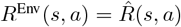 following *R*(*s, a*) = *R*^Bonus^(*s, a*) + *R*^Env^(*s, a*). In our simulations, we used 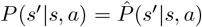 and *Q*(*s, a*) = *R*(*s, a*) = *R*^Bonus^(*s, a*), as there is no reward in the environment.

The curiosity models we present are independent of the learning technique used by the agent to learn *P* (*s, a*). Hence, the distribution can be learned using an alpha learning rate [49] or a Bayesian approach [64] instead of the rolling window *H* used in this article. A different choice for model learning may provide slightly different results.

To adapt our approach to multi-step environments, agents may plan on their model using dynamic programming techniques such as value-iteration. This consists in computing the expected utility *Q*(*s, a*) of each action *a* in state *s* based on the learned models *P* and *R*. The Q-values are computed using the Bellman update rule [39] following

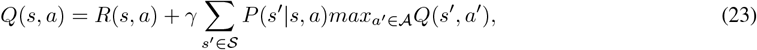

where *γ* ∈ [0, 1[is a discount factor. For one-step environment, the agent does not plan and directly uses *γ* = 0 and *Q*(*s, a*) = *R*(*s, a*).

## B Comparison of performance

### B.1 Distance to the true model of the environment

**Figure S1:**
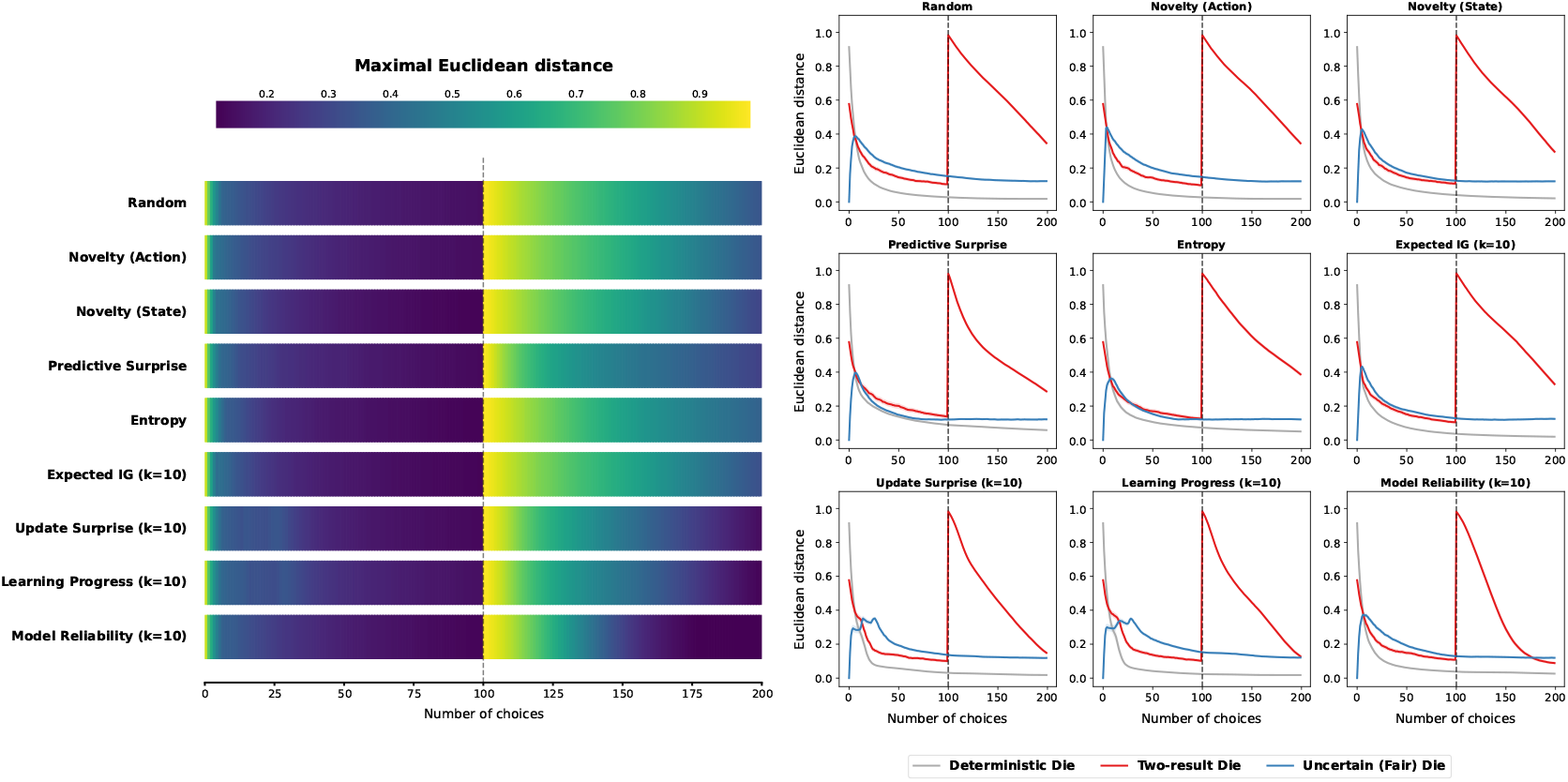
Evolution of the maximal Euclidean distance between the learned probability distributions and the ones from the environment over time. The darker the color, the best the agent understands the true underlying distributions.

Euclidean distance between the model learned by the agent and the transitions of the environment describes how well the agent understands the environment it interacts with (Figure S1). At start, the agent expects all dice to be fair: the distance to the distribution of the uncertain die is equal to 0 and is maximal with the deterministic die (Figure S1, Right). The evolution of distance values reflect the choice preferences described in the main article (Figure 3). In early trials, methods which focus on the uncertain die such as predictive surprise, entropy, or state novelty, approximate the distribution of the uncertain die better than other methods, but may take longer to learn the two other distributions (Figure S1, Right). The sequential learning induced by the update surprise and learning progress bonuses delays the acquisition of an accurate model for the uncertain die. The model reliability, the learning progress and the update surprise bonuses help the agent re-learn the distribution of the two-result die after the task change.

### B.2 Fastest change detection

To examine which information bonus allows the fastest adaptation to changes, we compare the information bonuses for the two-result die for model reliability, learning progress and update surprise (Figure S2). The information bonus for model reliability increases at least as fast as the two other methods when comparing the information bonuses directly (Figure S2, Left). Forcing the agents to have a random behavior at all steps shows that the information bonus for model reliability may even increase faster than the ones for update surprise and learning progress methods (Figure S2, Right). This indicates that model reliability is an efficient information bonus to direct exploration when trying to adapt to task changes.

**Figure S2:**
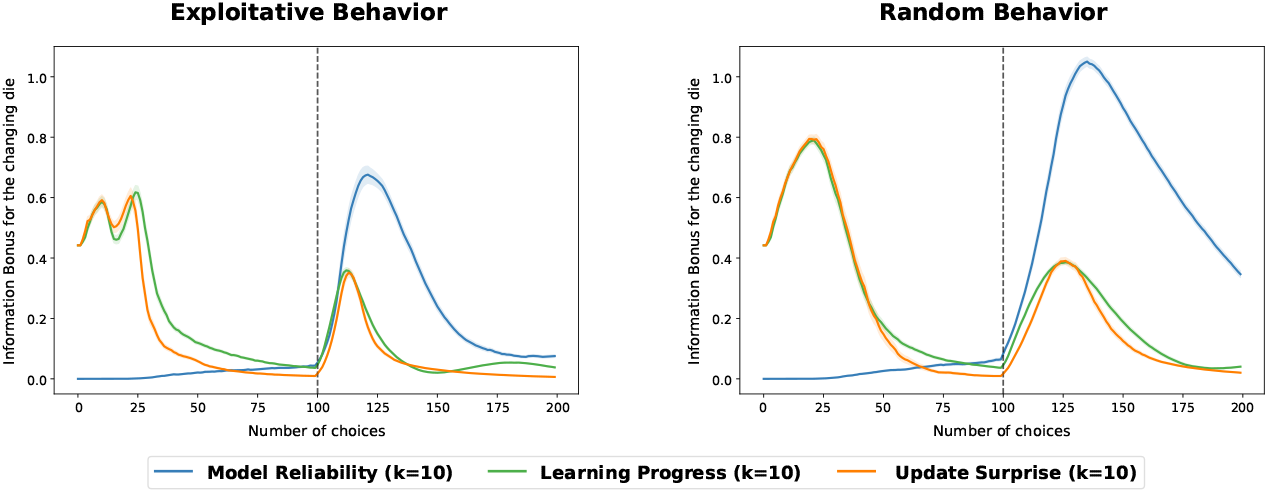
Evolution of the information bonus for the changing die over time for the model reliability, the learning progress and the update surprise methods. All agents use *k* = 10 and either use *β* = 10 (Left) or act randomly with *β* = 0 (Right). Each plot is generated over 500 seeds of 200 trials, with the two outcomes of the two-result die changing at step *t* = 100.

## C Additional simulations and results

### C.1 One-step information bonuses (k=1)

**Figure S3:**
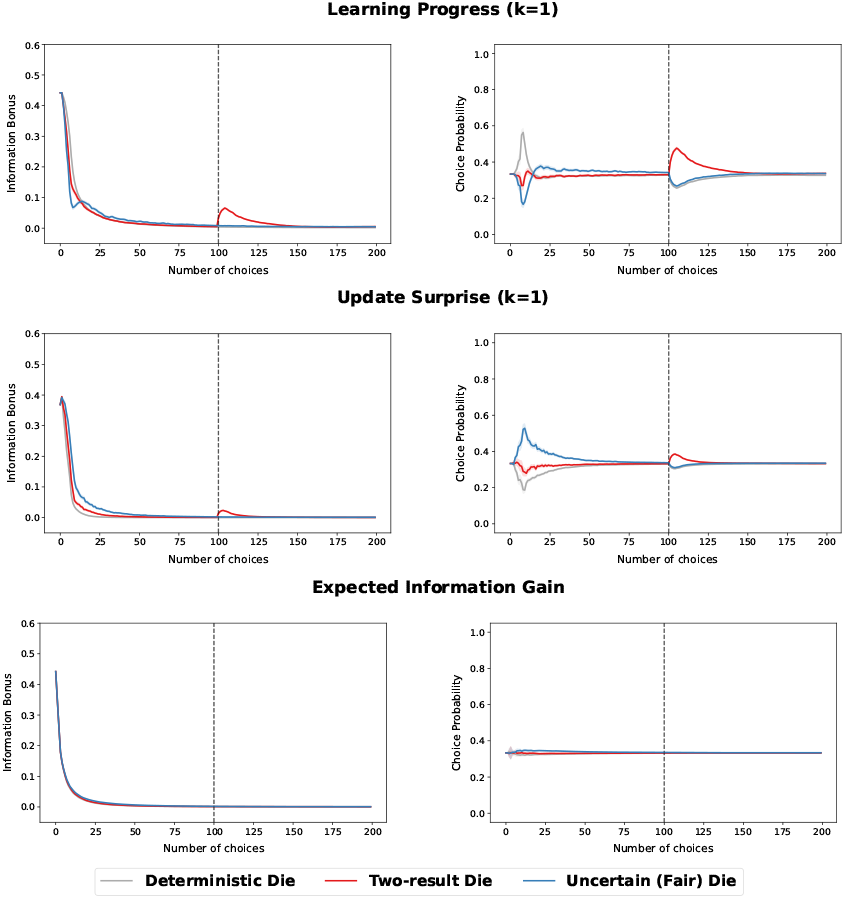
Evolution of information bonuses (Left) and choice preferences (Right) over time for the learning progress (Top), the update surprise (Center), and the expected information gain (Bottom). Each plot is generated over 500 seeds and 200 trials, with the two outcomes of the two-result die changing at step *t* = 100.

In the main article, we presented 10-step variations for learning progress, update surprise and expected information gain. For the sake of completeness and to understand each method better, we show the plots for 1-step variations (Figure S3). All information bonuses are lower with *k* = 1 than with *k* = 10 (Figure 2, Right) as the learned models vary less with one one observation than ten. The three bonuses generally display the same results as for *k* = 10, with decreasing information bonuses for all three methods and an increase when the task changes for learning progress and update surprise. As for *k* = 10, expected information gain shows a faint preference for the uncertain die in the stable environment. However, when the task changes, the information bonuses for expected information gain are too little to show an increase for the two-result die, as opposed to the results for *k* = 10. The main difference between the plots for *k* = 1 and *k* = 10 is that update surprise with *k* = 1 prefers the uncertain die in the early trials (Figure S3, Center) while the *k* = 10 variation prefers the deterministic and the two-result dice in early trials. This difference stems from the fact that two different observations induce a bigger second-step update than two similar observations. Hence, after two trials, the one-step update of the model for the deterministic die usually becomes smaller than for the uncertain die. On the contrary, the one-step decrease in entropy remains bigger for the deterministic die than the fair one for a few trials, due to the initial uniform prior. Thus, the learning progress results for *k* = 1 and *k* = 10 match, with an early preference for the deterministic and the two-result dice (Figure S3, Top).

### C.2 A bi-phasic information bonus for learning progress and update surprise

**Figure S4:**
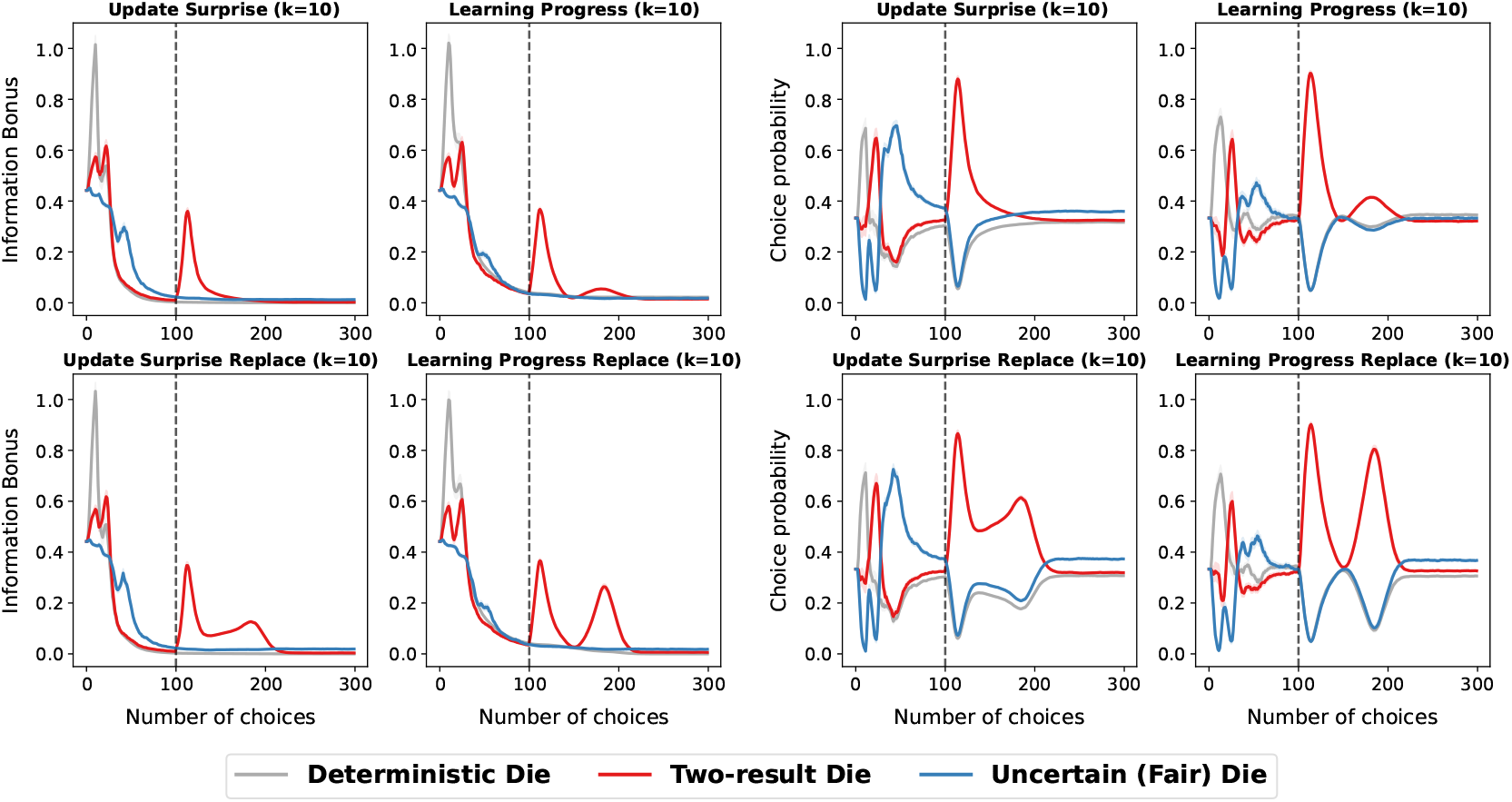
Evolution of information bonuses (Left columns) and choice preferences (Right columns) over time for update surprise and learning progress without (Top) or with (Bottom) replacement. Each plot is generated over 500 seeds and 300 trials, with the two outcomes of the two-result die changing at step *t* = 100. The plots for learning progress and update surprise without replacement correspond to the ones presented in the main article, with an extended time-window of *t* = 300 trials.

The learning progress and update surprise methods we presented use an horizon of *H* = 50, take the *k* = 10 most recent observations out of the model, and compare a distribution formed of *H* − *k* = 40 observations with the current distribution of *H* = 50 observations. Another potential implementation is to compare the current model with the actual model the agent was using *k* steps before. With this method, the *k* most recent observations are taken out and replaced with *k* older observations. Hence, we name this approach the *replace* variation.

The *replace* variation displays similar early information bonuses to the original plots. However, when facing the change in the two-result die distribution, the *replace* version of learning progress and update surprise show a bi-phasic information bonus (Figure S4). This phenomenon is particularly salient for learning progress computed as a difference of entropy, as learning progress for the two-result die reduces to 0 several trials after the task change, before increasing again. This bi-phasic evolution happens when two models have the same entropy although they are different, leading to a temporary null learning progress. In our example, since we use finite horizons of *H* = 50 and *k* = 10, the entropy of the two-result die increases from the one of a two-result uniform distribution to a four-result uniform distribution, and then decreases back to the entropy of a two-result uniform distribution. This increase and decrease in entropy leads to a null learning progress when the two learned models separated by *k* = 10 observations have the same entropy. This bi-phasic exploration can appear in a broad range of curiosity models computing variations, particularly when the method to learn models uses a rolling window.

### C.3 Predictive surprise and Entropy

**Figure S5:**
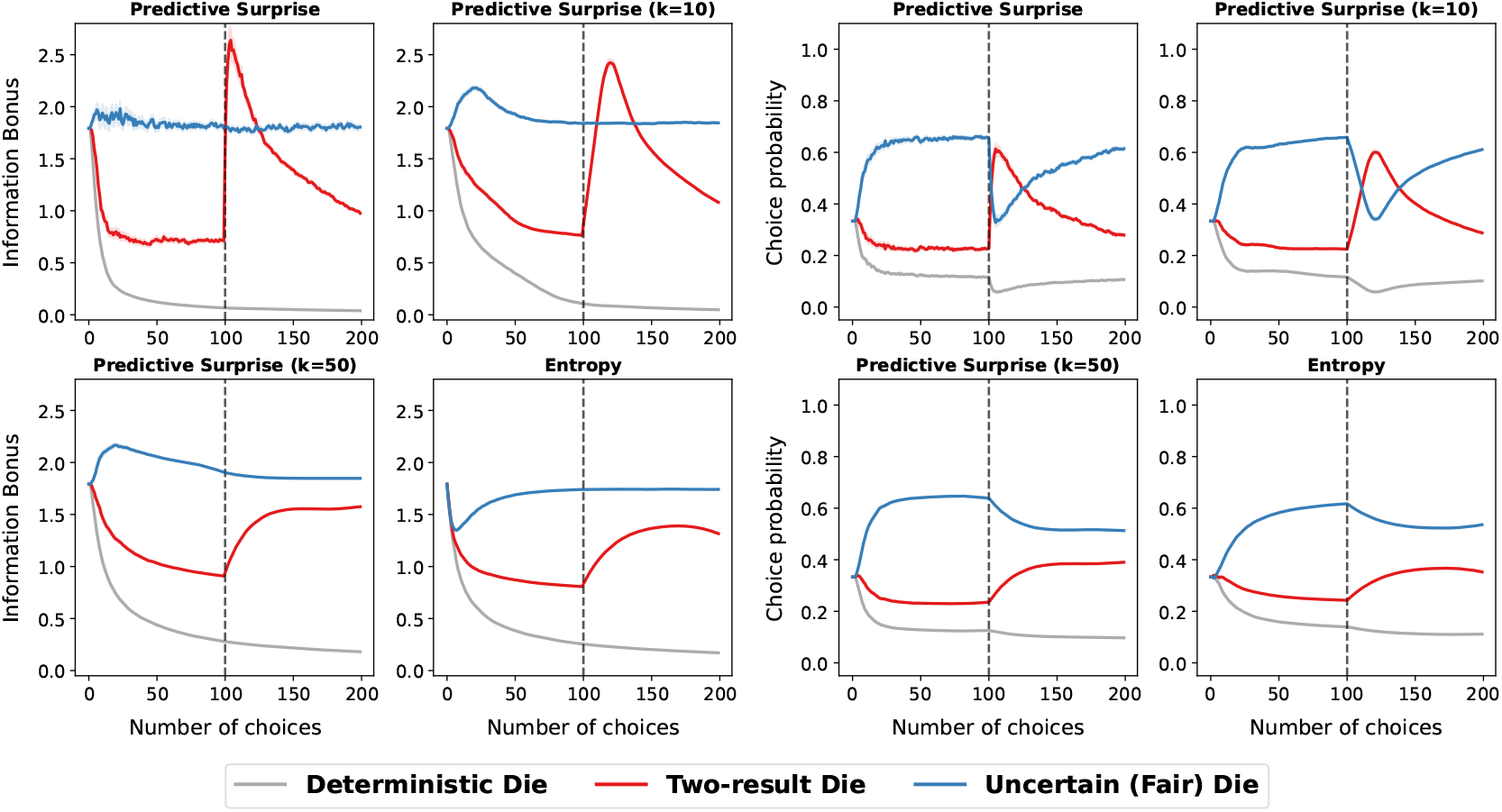
Evolution of information bonuses (Left columns) and choice preferences (Right columns) over time for predictive surprises and entropy. The plots show the results for immediate predictive surprise, predictive surprise averaged over the *k* = 10 or the *k* = 50 last observations, and entropy. Each plot is generated over 500 seeds and 200 trials, with the two outcomes of the two-result die changing at step *t* = 100.

In the main article, we normalized predictive surprise and entropy to compare the two information bonuses with other approaches. In this section, we do not normalize the information bonuses and increase exploration by lowering *β* from 3 to 1. Increasing exploration is mandatory to favor choice variability for both methods, as they have high information bonuses and can easily be attracted to uncertain attractors. We compare predictive surprise, averaged predictive surprise over the last *k* = 10 or *k* = 50 observations, and entropy (Figure S5). Similar to entropy, predictive surprise is attracted to irreducible uncertainty but varies more when not averaged over many observations. However, predictive surprise detects task changes better than entropy (especially for *k* = 1 and *k* = 10). Even when averaged over *k* = 50 observations, surprise and entropy do not match as entropy represents a prospective expected prediction, while surprise is retrospective and uses the model at the time of the observation. Averaging surprise over several trials favors stability but adds a delay between the observations and the update of the information bonus, which is particularly visible in early trials or at task change (Figure S5).

### C.4 Inverting the order of the distributions in update surprise

Many work invert the order of the prior and posterior for the computation of update surprise (section 2.2.2). However, this inversion reduces and delays the information bonus when facing surprising events (Figure S6, Bottom). This phenomenon stems from the asymmetry of the KL divergence. If the first distribution predicts a new observation while the second does not, the KL divergence diverges (division by 0). Conversely, if the second distribution predicts an observation that the first one does not predict, the KL divergence does not diverge and increases slightly (as the distribution probabilities become more dissimilar). Thus, in our simulations, putting the current distribution first and the old distribution second produces a larger surprise signal than doing the opposite (Figure S6). This effect would further increase without a uniform prior which guarantees that there is no division by 0.

**Figure S6:**
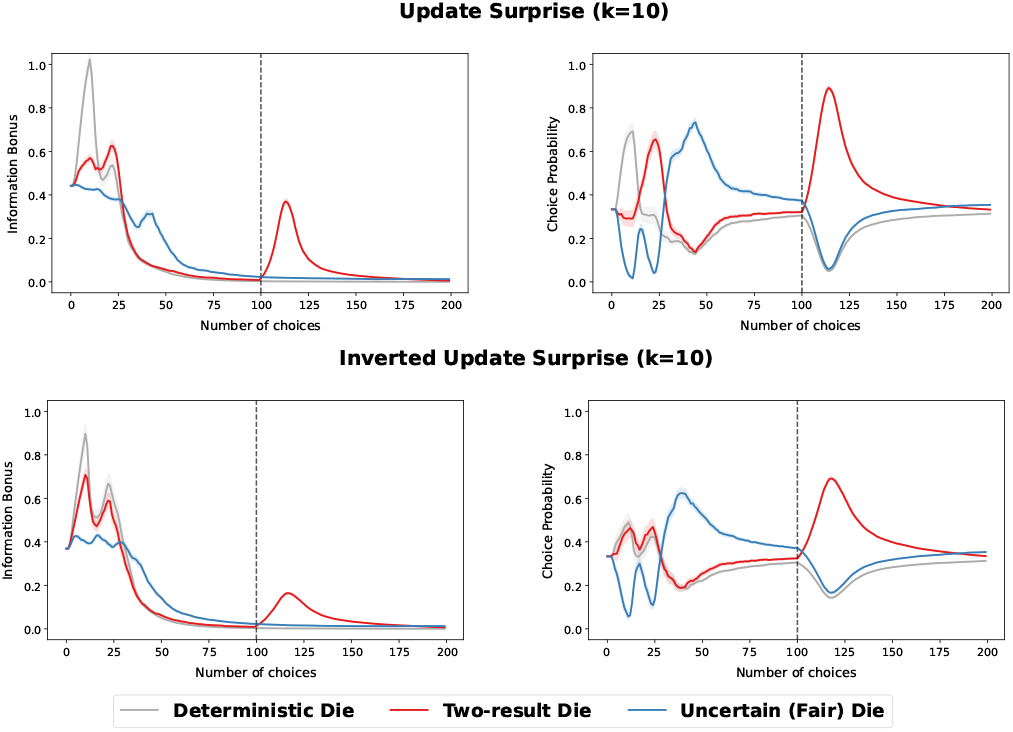
Evolution of information bonuses and choice preferences over time for the update surprise (Top) and the inverted update surprise (Bottom). Each plot is generated over 500 seeds and 200 trials, with the two outcomes of the two-result die changing at step *t* = 100.

### C.5 Why learning progress needs absolute values?

**Figure S7:**
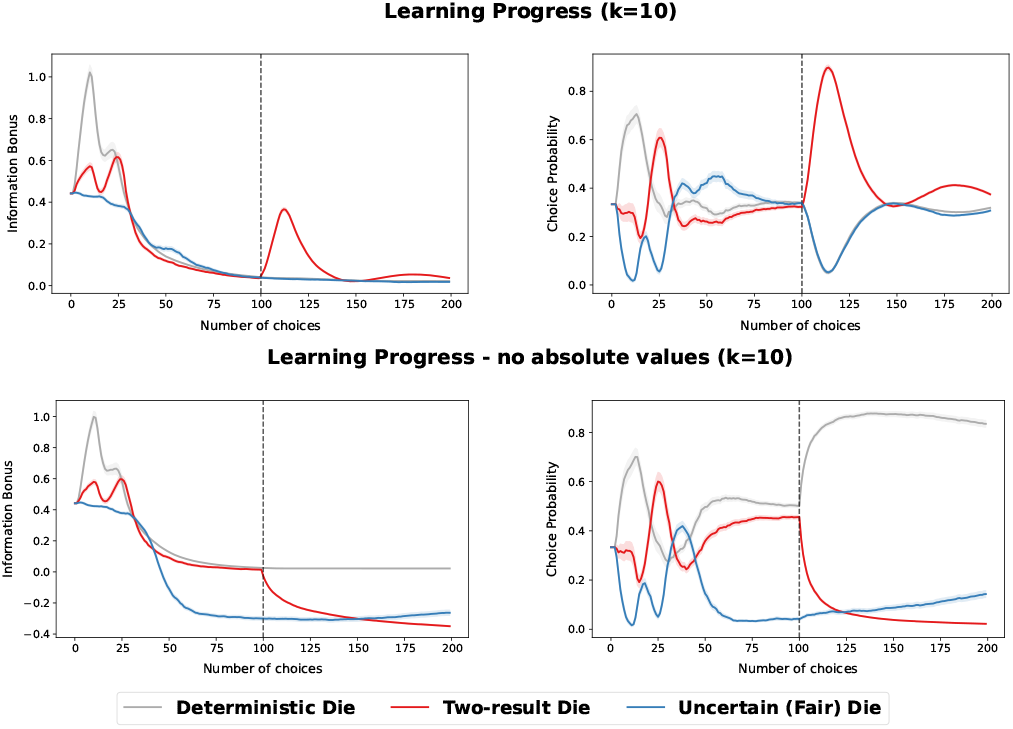
Evolution of information bonuses (Left) and choice preferences (Right) over time for the learning progress with (Top) and without (Bottom) absolute values. Each plot is generated over 500 seeds and 200 trials, with the two outcomes of the two-result die changing at step *t* = 100.

Originally, learning progress would only favor increases in the predictive power of the learning agent [73]. This equates to using the formula of learning progress we introduced without absolute values (Equation 9). Without absolute values, decreases in the predictive power of the agent lead to negative learning progress. Thus, agents which use this information bonus must avoid surprising events that do not improve their predictive power. Surprising events often happen for uncertain or changing actions. Thus, in our experiment, using learning progress without absolute values leads to the agent quickly avoiding the fair die (Figure S7, bottom). Similarly, when the two-result die changes, the entropy increases and the agent starts avoiding the two-result die as well. To avoid this phenomenon which leads to worsened learning and does not correspond to animal behavior, recent learning progress models use absolute values, making the hypothesis that the agent can learn as much from decreases in predictive power than increases [78]. Based on the task at hand, one can choose to make increases in the predictive power of the agent more important than decreases, or vice versa, by using an additional weighting parameter instead of absolute values.

### C.6 Random goal-generation

**Figure S8:**
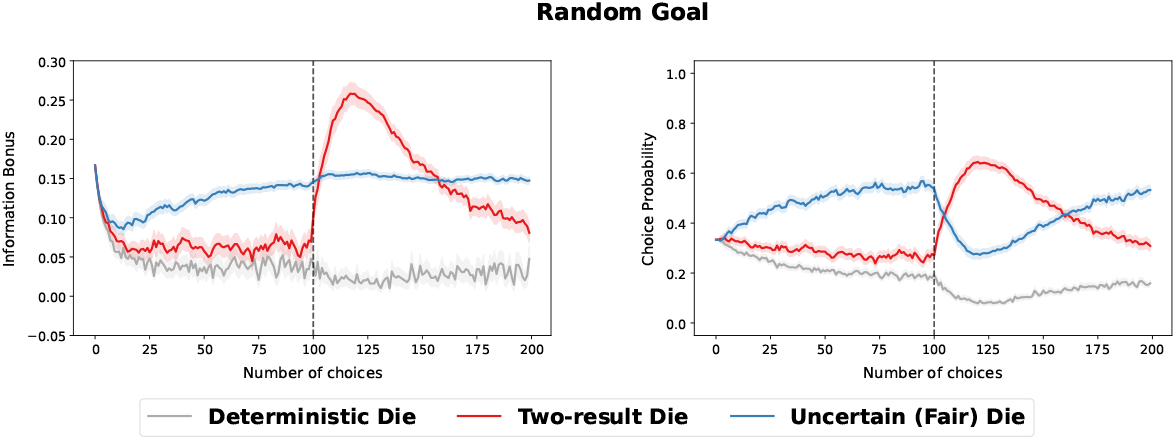
Evolution of information bonuses (Left) and choice preferences (Right) over time for a goal-based agent with random goal generation. Each plot is generated over 500 seeds and 200 trials, with the two outcomes of the two-result die changing at step *t* = 100.

We introduce a random goal-generation agent to model a simple goal-based approach. This agent samples a goal in the observational space randomly (1 to 6) and tries to achieve it: the current goal does not change until the agent achieves it. To motivate the agent to choose the actions which lead to the goal state, we use the learned transition probabilities to the goal as information bonuses. With *G* the current goal of the agent, 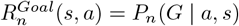. For example, if the goal of the agent is to obtain 1, the information bonus will be high for the deterministic die, low for the fair one and very low for the two-result one (respectively 1, 1*/*6, and 0 with perfect models). The goal-based method we introduce prefers the fair die in stable environments on average (Figure S8). This preference arises because the agent spends more time trying to achieve hard goals than easy ones. When the task changes, the agent prefers the two-result die over the fair die on average. This change in preferences mainly arises when the agent wants to attain 2 or 3 right after the change: the agent may roll the two-result die several time before re-learning the probability distribution (that now returns 4 and 5 with 50% chance each) and preferring the fair die for 2 and 3s. Overall, the results for the random goal generation are less general than the other ones presented, notably because they can be tweaked in many directions based on the parameters used or the goal-achievement method selected.

